# SARS-CoV-2 vaccination of laboratory rhesus monkeys (Macaca mulatta): Monitoring and efficacy

**DOI:** 10.1101/2022.11.11.516206

**Authors:** Dan Qi Priscilla Oh, Iris Grothe, Herbert Lukaß, Andreas K. Kreiter, Markus Hoffmann, Detlef Wegener

## Abstract

The availability of effective vaccines and a high vaccination rate allowed the recent mitigation, or even withdrawal, of many protective measures for containing the SARS CoV-2 pandemic. At the same time, new and highly mutated variants of the virus are found to have significantly higher transmissibility and reduced vaccine efficacy, thus causing high infection rates during the third year of the pandemic. The combination of reduced measures and increased infectivity poses a particular risk for unvaccinated individuals, including animals susceptible to the virus. Among the latter, non-human primates (NHPs) are particularly vulnerable. They serve as important models in various fields of biomedical research and because of their cognitive capabilities, they receive particular attention in animal welfare regulations around the world. Yet, although they played an extraordinarily important role for developing and testing vaccines against SARS-CoV-2, the protection of captive rhesus monkeys against Covid-19 has rarely been discussed. We here report upon twofold mRNA vaccination of a cohort of 19 rhesus monkeys (*Macaca mulatta*) against infection by SARS-CoV-2. All animals were closely monitored on possible side effects of vaccination, and were tested for neutralising antibodies against the virus. The data show that vaccination of rhesus monkeys is a safe and reliable measure to protect these animals against SARS-CoV-2.

## Introduction

In its third year, the SARS-CoV-2 pandemic has reached its highest incidences so far. Due to new, highly mutated variants of the virus with significantly higher transmissibility and reduced vaccine efficacy^1-4^, the number of cases increased from about 100 million in 2020 to 300 million in 2021, and to more than 600 million since (World Health Organization, Coronavirus Dashboard, https://covid19.who.int, accessed 13.09.2022). A high vaccination rate and the basic effectiveness of the available vaccines also against the new subtypes of the virus^5, 6^ prevented a corresponding increase in hospitalisations and admissions to intensive care and enabled the reduction of many protective measures against the virus.

For non-vaccinated individuals, as well as for animals susceptible to the virus^7, 8^, though, more liberal protective measures increase the risk of infection by the virus. Among susceptible animals, Old World monkeys (Catarrhini) are particularly vulnerable. They exhibit the same twelve key amino acid residues in the Angiotensin-converting enzyme 2 (ACE2) receptor, the contact surface to enter host cells, as humans^9, 10^.

Among the Catarrhini, rhesus macaques (*Macaca mulatta*), cynomolgus macaques (*Macaca fascicularis*), and the African Green Monkey (*Chlorocebus sabaeus*) are easily infected by the virus and exhibit symptoms of disease similar to those in humans. They have become important models for research on the physiology and treatment of the disease, including the neurological symptoms associated with the so-called “long COVID”^11-19^. Rhesus monkeys, in particular, are capable of displaying severe symptoms after infection. They served in pre-clinical testing for each of the five vaccines now approved for human use by European and/or U.S. regulatory agencies^20-24^.

Despite their role in the development and testing of SARS-CoV-2 vaccines and the treatment of COVID-19, little details about possible side effects following vaccination of rhesus monkeys have been reported. Because they serve as important models for a variety of biomedical research fields^25-30^, many animals live in research facilities. Even in these husbandries, despite their high hygiene and biosafety requirements, complete protection against pathogens from the environment is usually not possible. The likelihood of contact with SARS-CoV-2 will increase as general societal protective measures become less stringent. Therefore, measures for the health and wellbeing of these research animals should now, under the current conditions of the pandemic, include their protection against SARS-CoV-2 infection.

We here report the vaccination of a cohort of 19 rhesus macaques with the BNT162b2 vaccine (BioNTech/Pfizer), including thorough pre- and post-vaccination investigation, with special emphasis on body temperature and body weight. Vaccination was performed due to veterinary advice. Our report is thought to serve as a reference for research institutes and zoos currently reasoning the pros and cons of vaccinating their animals.

## Methods

Animal housing, behavioural, and veterinary procedures followed the Regulation for the Welfare of Experimental Animals issued by the Federal Government of Germany and were approved by the local authorities. All monkeys (*N* = 19) were bred for scientific purposes and were obtained from the German Primate Centre (DPZ, Göttingen). They were housed in large, environmentally enriched compartments, in groups of two or four monkeys. Three individuals were single-housed, with visual and auditory contact with other monkeys. A detailed description of the housing conditions can be found elsewhere^31^.

Monkeys were aged between 9 and 20 years at the time of first vaccination. Body weights were between 8.4 and 18.2 kg. All monkeys were healthy and in good overall condition. They participated in neuroscience laboratory procedures, including periods of restricted water access^31^, yet all received full food and water supply from at least three days before vaccination to five days after vaccination. Monkeys were vaccinated using the BNT162b2 (BioNTech/Pfizer) vaccine^23^. Following the instructions from the supplier, the vaccine concentrate was diluted with 1.3 ml of a 0.9% sodium chloride injection. Each monkey was injected with 0.2 ml (equivalent to a dose of 10 μg) intramuscularly.

All procedures were done in awake monkeys. The monkeys entered a horizontal primate chair, in which they sat in a Sphinx-like position. The rear of the chair was opened for administering the vaccine and conducting follow-up examinations. Vaccination was scheduled in two groups - ten monkeys in the first and nine in the second. All monkeys received a second injection six weeks after the first vaccination.

Before vaccination, body weight and rectal temperature were taken from all monkeys. The follow-ups were carried out for seven days after vaccination at approximately the same time of the day and included daily control of body temperature and body weight as well as inspection of injection sites and checking for cold symptoms (runny nose, tearing eyes, sneezing, coughing) and appetite. If body temperature, general condition, and overall behaviour were unremarkable, measuring of temperature and weight were suspended at the weekends.

Three animals were tested for neutralising antibodies against SARS-CoV-2 prior to vaccination and four to five weeks after first and second vaccination. All other animals were tested after second vaccination only. The test used pseudovirus particles^32^ equipped with the spike (S) protein of either the early B.1 lineage, or the currently dominating BA.4 and BA.5 lineages (identical on amino acid level). For this study, we used a replication-deficient vesicular stomatitis virus coding for eGFP and firefly luciferase (VSV*ΔG-FLuc^33^, kindly provided by Gert Zimmer, Institute of Virology and Immunology, Mittelhäusern and Bern, Switzerland). Pseudovirus particles were produced according to an established protocol^34^, incubated for 30 min at 37 °C with serial dilutions of heat-inactivated blood serum and then added to target cells (Vero cell line). A detailed protocol of the pseudovirus neutralisation assay can be found elsewhere^35^. If the serum contains neutralising antibodies against the S protein embedded in the pseudovirus particle, they will bind to the S protein and block pseudovirus cell entry. As a result, the number of infected cells will decrease depending on the concentration of neutralising antibodies in the sample, resulting in reduced activity of pseudovirus-encoded firefly luciferase in cell lysates, which serves as a surrogate for pseudovirus cell entry. Finally, the serum dilution that leads to half-maximal inhibition of pseudovirus cell entry (NT50, neutralising titer 50%) was determined. The higher the NT50, the stronger is the neutralising activity of the sample.

NT50 was calculated by estimating the relative inhibition of pseudovirus cell entry at ascending dilution factors of the serum. The NT50 is equivalent to the IC 50 obtained from a nonlinear fit to the dose-response curve. Data were analysed and visualised using GraphPad Prism (Dotmatics, Boston, MA) and Matlab (The MathWorks, Natick, MA). If not stated otherwise, statistical testing was performed using the Wilcoxon signed rank test, at 95% significance level. Effect size was calculated as Cohen’s d, using the standard deviation of pre-test data as reference.

## Results

Nineteen rhesus monkeys, subdivided into two cohorts of ten and nine monkeys, received two doses of the BNT162b2 (BioNTech/Pfizer) vaccine, diluted to the concentration recommended for children. The second cohort was vaccinated three weeks after the first, and the interval between the two vaccinations was six weeks. All animals were tested for body temperature and body weight on the day of vaccination and during seven days thereafter. The injection site was closely inspected whenever body temperature was taken, and all animals were daily monitored for their appetite, behaviour, cold symptoms, and general appearance. Blood serum of three example animals was used to test and quantify the presence of antibodies to the S proteins of SARS-CoV-2 lineages B.1 and BA.4/BA.5 before and after the first vaccination, and the serum of all animals was tested after the second vaccination.

### General appearance and overall behavior

Monitoring of monkeys by animal caretakers and scientific staff (DW, DQPO, IG) did not provide evidence of significant physiological effects during the seven days following vaccination. No signs of cold symptoms and no changes in general appearance were detected and all monkeys seemed to have a normal appetite. Injection sites were unremarkable - daily inspection did not reveal redness or hardening, and monkeys did not show specific responses to palpation. A few deviations were noted at the behavioural level: One monkey revealed a slight change in motion profile and sometimes preferred climbing onto a platform, rope, or other housing element instead of jumping. This change in behaviour was observed after both injections and lasted for two days. Two other monkeys were described to appear a bit tired during a few days after first vaccination, and two other monkeys were described to be significantly less active than usual during a period of three to four days in-between the two vaccinations, about 2.5 weeks after the first vaccination.

### Body temperature and body weight

Figure 1 summarises the data on body temperature and body weight during the week after first vaccination, relative to temperature and weight at the day of vaccination. Because no animal showed a consistent sign of reduced well-being, measurements were suspended at the weekend and were performed during days 1, 2, 3, 6, and 7 after vaccination, providing a total of 95 values each for body temperature and body weight. For body temperature, 22 values deviated by more than 0.5°C from the value at the day of vaccination, and half of those (12 %) were higher than the initial temperature (Fig. 1A). Exactly one week after vaccination, the body temperature of five animals deviated by more than 0.5° from the initial temperature. It was higher in two animals and it was lower in three other animals as compared to the day of vaccination. There was no statistical indication in the group of animals for a consistent change of body temperature one week after vaccination following paired testing (*Z*(1,18) = 0.262, *P* = 0.79) (Fig. 1B). For body weight, 23 values deviated by more than 1% of the body weight at the day of vaccination, and 21 of those (22%) were indicating a loss of weight (Fig. 1C). One week after vaccination, 1 monkey gained more than 1% of body weight, while six animals lost more than 1%. Though hardly recognisable by observation of behaviour and overall appetite, this constitutes a significant effect on body weight (*Z*(1,18) = 2.399, *P* = 0.016) (Fig. 1D). Effect size, however, was negligible (*d* = 0.0352) and there was no relation to body temperature (Pearson correlation, *P* = 0.66) (Fig. 1E).

**Figure 1.**
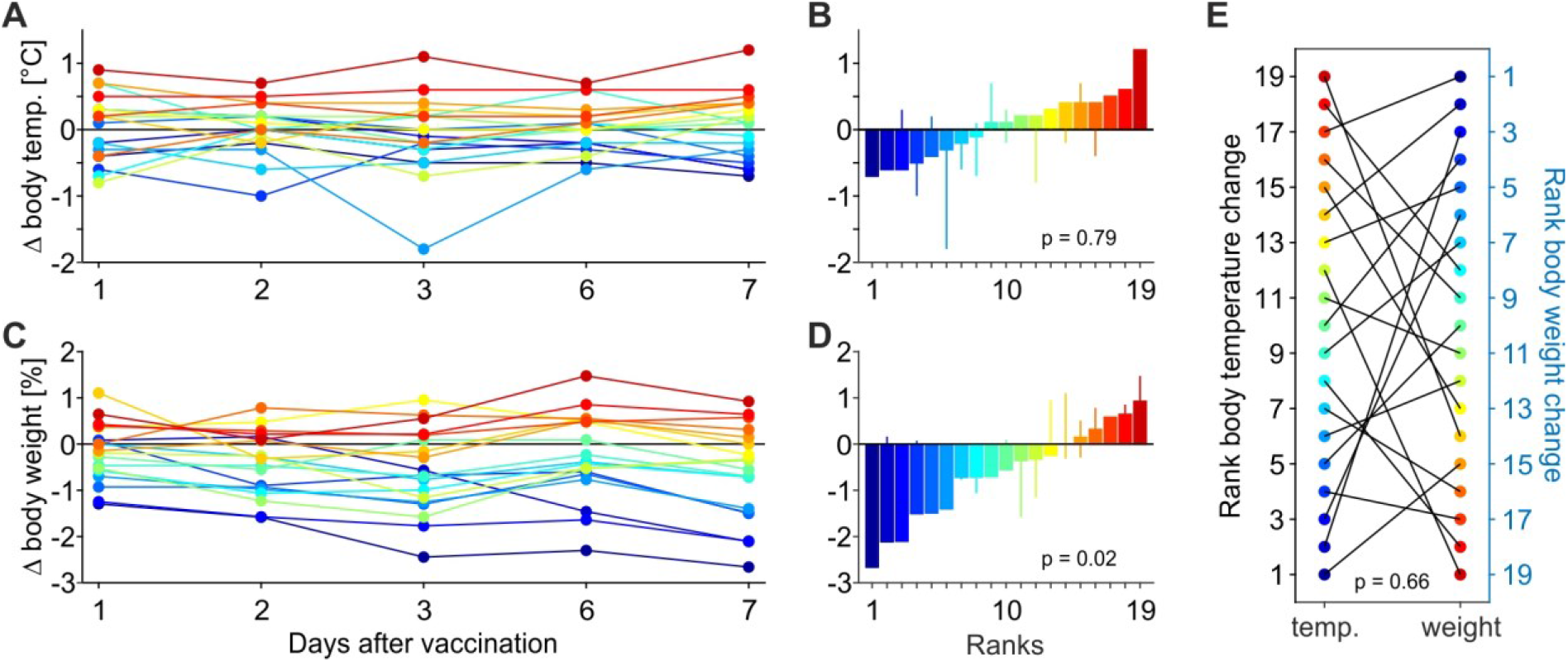
Deviations of body temperature and body weight after first vaccination. (**A**) Body temperature of *N* = 19 monkeys at five days during the week following vaccination. Body temperature taken at day of vaccination serves as reference (black horizontal line). (**B**) Body temperature deviation sorted by magnitude. Bars indicate body temperature deviation one week after vaccination, thin lines indicate minima and maxima during the week of measurement. (**C, D**) Body weight deviation during the week following the vaccination. Conventions as in (A, B). (**E**) Correlation of body temperature (ascending) and body weight (descending) ranks. Black lines connect the ranks of individual animals.

Figure 2 shows the corresponding measurements obtained during the week following the second vaccination, again referenced to the day of vaccination. Data were similar to the results obtained after the first vaccination. Body temperature was unremarkable (*Z*(1,18) = 0, *P* = 1), slight increases in temperature were as frequent as slight decreases (Fig. 2A, B). Body weight changes were again significant (*Z*(1,18) = 3.019, *P* = 0.003) with nine animals having a weight loss of more than 1% (Fig. 2C, D). Effect size was still negligible (*d* = 0.0598) and the weight loss did not correlate with temperature (Pearson correlation, *P* = 0.49) (Fig. 2E). Yet, one individual revealed a loss of almost 5% one week after vaccination, increasing to almost 17% about three months after second vaccination. The weight then stabilized. About 2.5 weeks after the first vaccination, the monkey showed slightly reduced activity for a few days, but otherwise behaved inconspicuously and did not show any abnormalities during several veterinary checks.

**Figure 2.**
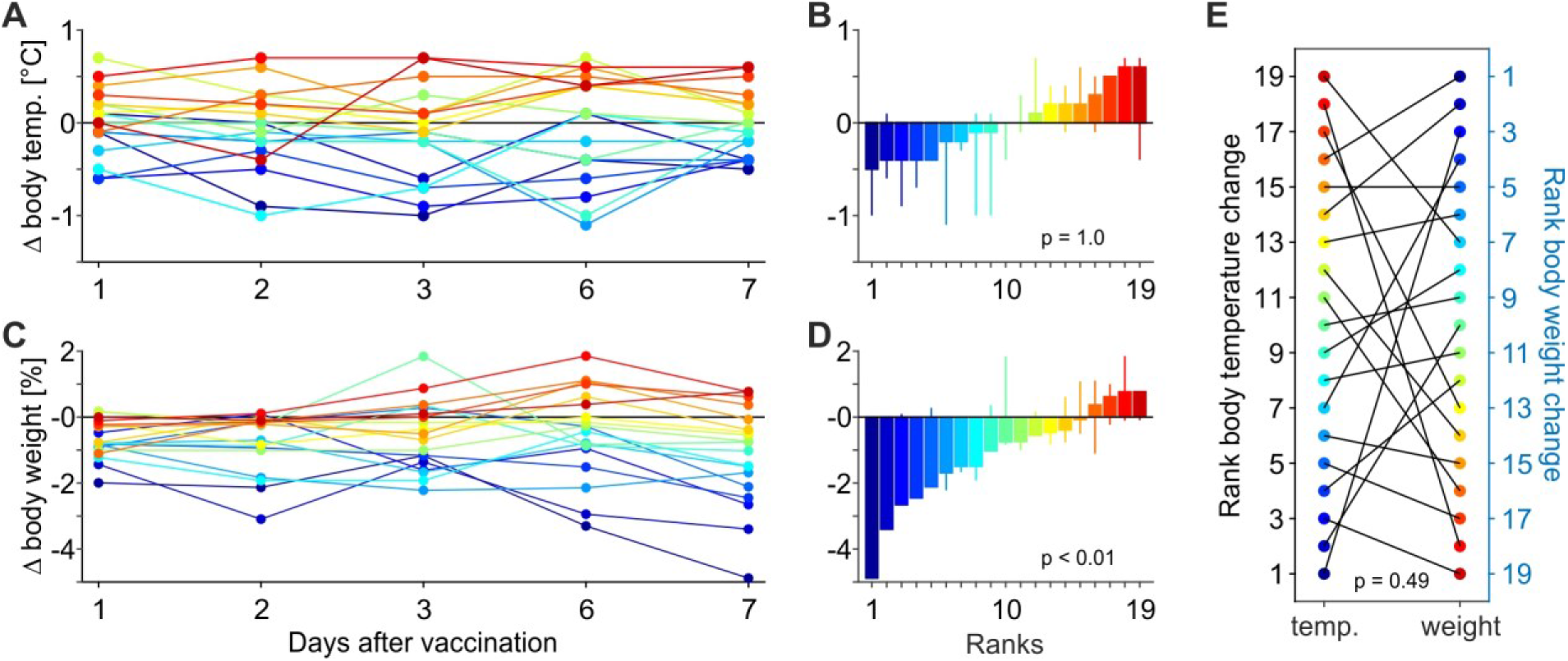
Deviations of body temperature and body weight after second vaccination. Conventions as in Fig. 1.

### Antibody testing

Presence of neutralizing antibodies against the SARS-CoV-2 S protein was investigated by a pseudovirus neutralisation test. The test was performed on blood serum from three example animals before and after first vaccination, and on serum from all animals after second vaccination. Pseudovirus particles were equipped with S proteins of different SARS-CoV-2 lineages to test the efficiency of the immune response against the B.1 lineage, which dominated the early phase of the pandemic, and against the BA.4/BA.5 lineages, which were dominating at the time of vaccination. No neutralising antibodies were detected in the serum of any of the three example animals before vaccination, but all were tested positive for neutralising antibodies after first vaccination (Fig. 3A). The blood serum of all but two animals (N = 19) showed significant neutralising activity after the second vaccination (Fig. 3B). Compared to human reference data^36, 37^ (*N* = 23), the neutralising antibody activity of the cohort of 19 macaques was weaker (Wilcoxon rank sum test, *Z* = 2.63, *P* = 0.0086, two-tailed) (Fig. 3C). Neutralising antibodies against the variants BA.4/BA.5 were basically absent in the three example animals after first vaccination. After the second vaccination, two of the example animals and 6 animals in the entire cohort were showing minor neutralising activity for the BA.4/BA.5 subtypes. The immune response did not differ between macaques and humans (*Z* = 0.39, *P* = 0.6935) (Fig. 3D – F).

**Figure 3.**
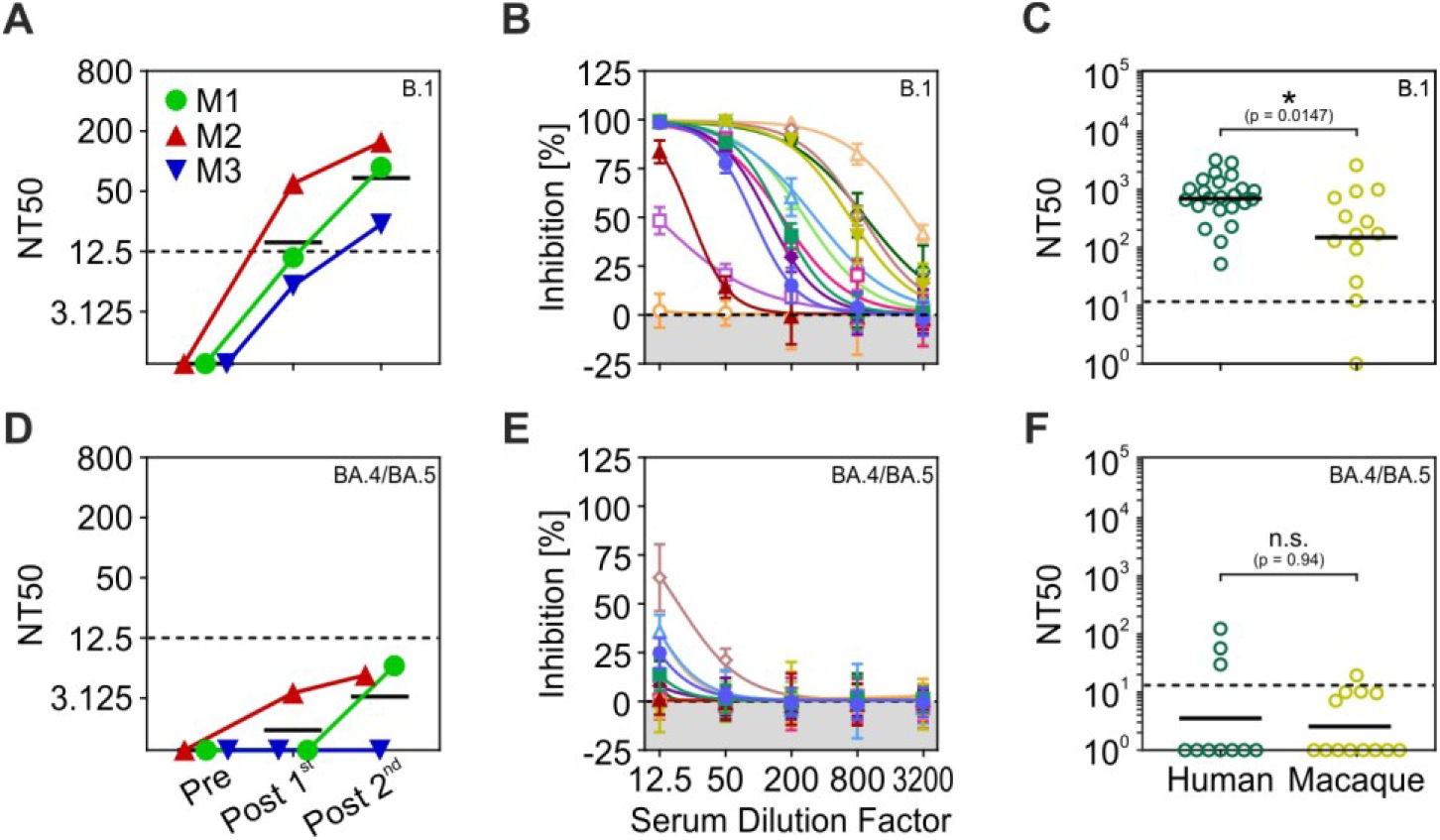
Blood serum neutralising activity against pseudoviruses bearing either B.1 or BA.4/BA.5 S proteins. (**A**) Neutralising activity (NT 50) of serum from three example monkeys against the SARS-CoV-2 B.1 lineage before (Pre), after first (Post1^st^), and after second (Post2^nd^) vaccination. (**B**) Inhibition of pseudovirus cell entry at different serum dilution factors for the SARS-CoV-2 B.1 lineage. Symbols indicate mean values of four repeated measurements, error bars indicate SD. Lines indicate nonlinear fit to data points for determining the titer at 50% neutralising activity (NT50). Each colour represents one subject. (**C**) Serum NT50 for the SARS-CoV-2 B.1 lineage in macaques compared to human reference data^36,37^. (**D** – **F**) Serum neutralising activity for the SARS-CoV-2 BA.4/BA.5 lineage. Conventions as in (A - C). NT 50 values below 6.25 were considered negative and were manually set as 1 in (C and F).

## Discussion

We here report the vaccination of laboratory rhesus monkeys with an adjusted dose of the BNT162b2 (BioNTech/Pfizer) vaccine, as recommended for children. The vaccine was developed based on the S protein sequence of the SARS-CoV-2 lineage dominating the early phase of the pandemic. Like other vaccines approved for human use, it was tested in rhesus monkeys before running clinical trials^23^. As of August 2022, the Strategic Advisory Group of Experts on Immunization (SAGE) of the World Health Organization considers the vaccine safe and effective and recommends its use for all individuals aged 6 months and above.

The interacting residues of the SARS-CoV-2 receptor, ACE2, are identical between rhesus monkeys and humans, making them similarly susceptible to the virus^8, 9^. A recent meta-analysis reports that of 129 rhesus monkeys exposed to the virus, all but two became infected^14^. Rhesus monkeys develop mild to moderate, occasionally severe respiratory disease after infection with SARS-CoV-2, lasting between 8 and 16 days. Tachypnea, dyspnea, asthenia, hunched posture, piloerection, reduced appetite, and pale appearance are among the symptoms of infection^11, 13, 17, 38-41^. Increases in body temperature as early as one day after infection^11, 12, 17, 38, 42^ and loss of body weight by up to 10% and more about 1 week after infection^11-13, 17, 40, 42^ have frequently been reported. Virus load is higher in older (> 15 years) animals than in younger ones. Additionally, immune response in older monkeys is delayed and more likely accompanied by a severe cytokine storm, persistent infiltration of lung tissue, and pro-inflammatory responses. Severe interstitial pneumonia is more frequent^13, 42-44^. Notably, swabs from the nose and throat were found to have high viral loads independent of the severity of the disease^11^. This increases the risk of cross-infection, especially when animals are not single-housed but kept in social groups.

The responsibility of maintaining the health and well-being of animals kept for scientific purposes calls for measures to protect them against infection by SARS-CoV-2. The primary measure to ensure this is an effective hygiene measurement and strict access control to the husbandry. Yet, SARS-CoV-2 variants with significantly increased transmissibility (like delta and omicron^45, 46^), evidence of high viral load in vaccinated humans in the absence of disease symptoms^5, 47^, and high incidences suggest an increased risk of the virus gaining entry into even secured husbandries. Vaccination of laboratory rhesus monkeys against SARS-CoV-2 to prevent (severe) disease hence mitigates the risk of health detoriation^20-24^ and prevents temporary or permanent exclusion of the animal from of the experiment. Thereby vaccination promotes laboratory animal welfare and prevents the need to replace any excluded animals. Although existing vaccines have limited efficiency against new emerging SARS-CoV-2 variants of concern, they are effectively preventing severe symptoms and hospitalization in humans, even over prolonged time^6, 48^.

We here show that after two injections of an adjusted dose of BNT162b2 (BioNTech/Pfizer), all but two rhesus monkeys have significant neutralising activity against the SARS-CoV-2 B.1 lineage, the early pandemic variant against which the vaccine was originally developed. Additionally, the blood serum of six monkeys had minor neutralising activity to the currently dominating SARS-CoV-2 BA.4/BA.5 lineages. This finding mirrors the situation in humans, where a third dose of the vaccine is needed to induce robust neutralising activity against BA.4/BA.5^36^. However, despite the ability of BA.4/BA.5 to efficiently evade antibody-mediated neutralisation, individuals vaccinated with two doses of BNT162b2 showed some neutralising activity against BA.4/BA.5^49^, even though the vaccine was developed against an earlier variant of the virus and BA.4/BA.5 possesses high potential for immune escape. The neutralising activity for BA.4/BA.5 in macaques was statistically not different from that of humans, but it was significantly weaker for the B.1 lineage. This might be due firstly to a generally weaker response of the rhesus immune system against the SARS-CoV-2 virus or against the antigen of the S protein encoded by the mRNA of the vaccine, secondly to a generally weaker stimulation of the rhesus immune system by the vaccine, or thirdly to a weaker immune response due to the lower dose of the vaccine, which was adjusted as recommended for children.

The observations reported in this paper have a few limitations. First, neutralisation was studied using a pseudovirus neutralisation assay. Although, this assay was shown to represent a suitable model to investigate SARS-CoV-2 cell entry and its neutralisation^32^, our data await formal confirmation with clinical SARS-CoV-2 isolates. Second, we did not investigate changes in neutralising activity over an extended period. As a consequence, we cannot make any statement on the longevity of neutralising antibody responses in rhesus macaques. Finally, we did not investigate T-cell mediated immune responses in rhesus macaques.

The vaccination had minor side effects, which are reported here to inform veterinarians and personnel. We were unable to detect visual abnormalities such as swelling or redness or specific responses of the monkeys to palpation of the injection site. Body temperatures fluctuated to some degree, yet we did not observe consistent changes in the group of monkeys and we did not identify changes at the level of individuals consistent with other signs of reduced well-being or weakness. We measured a loss of body weight over the group of animals after both the first and the second vaccination. Weight loss was statistically significant but the effect size was negligible. On the individual level, we identified one animal that was both losing more weight than others and also showing more weakness. Weakness stayed for only a few days but the weight loss increased over a time period of three months after vaccination before it eventually stabilized. The monkey was examined several times by a veterinarian, but the external appearance, blood and faecal samples, and imaging data revealed no physiological or other abnormalities, and the monkey behaved normally, including feeding behaviour. As such, we do not know whether the weight loss was triggered by the vaccination or by another factor.

We conclude that rhesus monkeys show minor side effects as a response to vaccination with an adjusted dose of the BNT162b2 vaccine. The antibody response is significantly weaker than in humans, most likely due to the smaller vaccination dose. Just as for humans, a booster vaccination is recommended to increase neutralising activity against the highly antibody-evasive omicron variant.

